# Repellent olfactory cues influence the optomotor response modulation in *Drosophila melanogaster*

**DOI:** 10.64898/2026.02.02.702750

**Authors:** Giulio Maria Menti, Matteo Bruzzone, Sara Zerbinati, Mauro Agostino Zordan, Patrizia Visentin, Andrea Drago, Marco Dal Maschio, Aram Megighian

**Author notes:** First co-authorship (Giulio Maria Menti, Matteo Bruzzone). Last authors (Marco Dal Maschio, Aram Megighian).

## Abstract

Animals need to precisely perceive and integrate the environmental cues to orient and select the appropriate motor responses required for navigating. This is the case, for instance, of the optokinetic reflex (OKR) and the optomotor response (OMR) in *Drosophila melanogaster*, where optic flow stimulation modulates the head or the body and legs motor activity respectively. Despite large bodies of literature on both the OKR and the OMR, there is still a limited understanding, in flies, of the impact on these responses of concomitant, and potentially conflicting, sensory inputs. To investigate this aspect, we used fruit flies walking on a sphere, presented with optic flow stimulation leading to the OMR together with the simultaneous exposure to olfactory stimulation, either using established repellent or masking compounds. We analysed the effect of different substances, and of their concentration, on the dynamics of the flies’ response to moving gratings, evaluating the fly walking path as well as average speed and duration. This analysis revealed several alterations between the compounds tested, in agreement with reported data on the simpler OKR. In conclusion, we show that concomitant exposure to repellents and maskers may consistently affect fundamental processes (the OKR and OMR) available to insects for informing themselves while navigating through the environment.

## Introduction

The navigational abilities of insects are remarkably close, if not equal, to the skills exhibited by vertebrates. Besides, navigational skills are fundamental for everyday activities, like searching for food, escaping from hazards, and social interactions. Unravelling the components and processes which generate and influence the repertoire of behaviours adopted by insects when exploring and moving in the environment is becoming of even more importance for tackling the challenges placed by some species to agriculture and human health.

In Europe, the increasing presence of alien insect species, exacerbated by the important climate variations, is posing a serious threat to cultivations, as is the case for the dipteran *D. suzukii* ^[1]^, which seriously threatens soft summer fruit cultivations. Likewise, public health is endangered by several disease-transmitting insect vectors such as mosquitoes and ticks which are expanding their geographical areas ^[2]^. Moreover, the 2012 EU Biocidal Product Regulation, for the sake of the population’s safety and environment preservation, shortened the list of compounds available for companies to distribute and consumers to purchase ^[3]^. A deeper comprehension of the effects of the repellent compounds publicly available for production and commercialization would increase the efficacy of repellent products and enhance global protection against pests. This is particularly important for the threat posed by mosquitoes, which can become vectors of several relevant diseases ^[4]^. Indeed, an efficient approach to study the problem could rely on the use of a more readily manageable and well-known model organism, such as the fruit fly *Drosophila melanogaster*. Drosophila is more easily reared and managed in the laboratory compared to mosquitoes, allows for sophisticated genetic manipulation, as well as deep behavioural and neurophysiological analysis; moreover, it shares the same brain organisation ^[5,6]^ and sensitivity to some known mosquito repellents ^[7,8]^. In fact, reports in these species confirm said olfactory sensitivity to a series of compounds, both natural - eugenol (**E**), lemongrass (**L**) - and synthetic - picaridin (**P**), IR3535^®^ (**I**) - independently from their ecological differences ^[7,8]^.

Recently there’s been renewed interest and application of new protocols for studying OMR responses in mosquitoes ^[9,10]^, when, in the past, studies on the mosquito’s visual system were fewer and constrained on visually guided behaviours ^[11]^. Given the similar response to visual and olfactory stimuli between mosquitoes and fruit flies, we thus opted for investigating the possible influence of repellents on visually guided behaviours in *D. melanogaster* by evaluating the multimodal sensory integration of vision and olfaction with respect to the OMR outcome.

To this end, we set up a behavioural paradigm similar to others described in literature ^[12,13]^ where adult female *Drosophila* were tethered at their back and with their legs walking on the surface of a sphere in response to moving grating visual stimuli ^[14–16]^. The paradigm envisages a temporal segment where visual stimulation is paired with the administration of different odorous repellent compounds (Fig. 1). We evaluated ten different conditions with both repellents (eugenol and lemongrass) and maskers (picaridin and IR3535^®^) delivered at two volume concentrations of **0.5**% and **1**% in mineral oil, which was used alone in the control condition.

**Figure 1.**
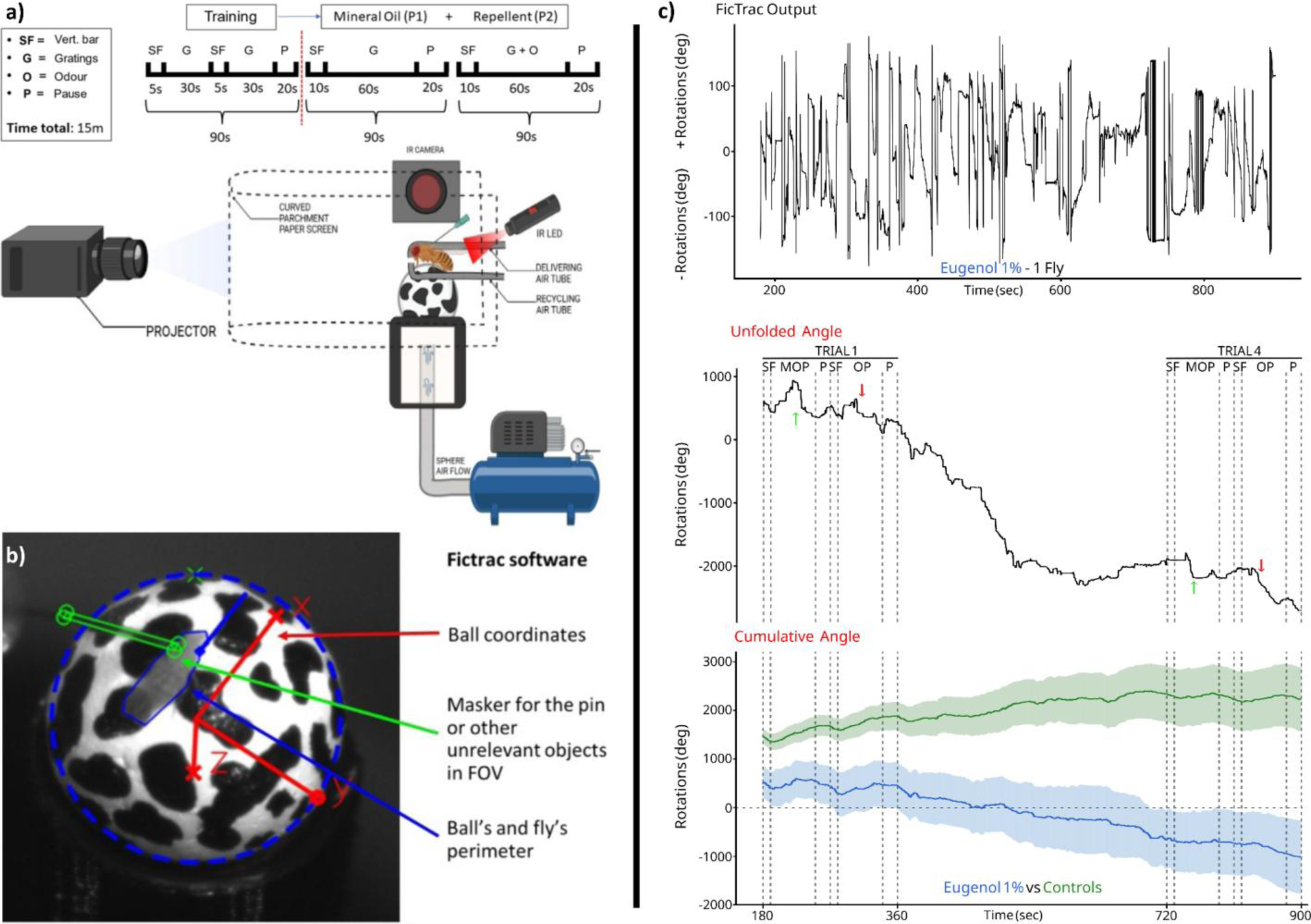
Methods and paradigm. (**a**) Experimental paradigm sequence and cartoon of the experimental apparatus. Created in BioRender. (**b**) FicTrac ^[41]^ GUI before processing one of our recordings. Blue ROIs identify the limits of the sphere and of the fly. Green ROIs mask object irrelevant or interfering with the tracking. Red arrows set the X, Y, Z axis allowing for the correct extraction of the space coordinates. (**c**) In descending order: example of one FicTrac ^[41]^ raw output, deconvoluted data, and one sample of the cumulated tracks from Controls (green) and Eugenol 1% (blue) groups (mean ± s.d.). **SF**: stripe fixation. **MOP**: mineral oil phase (plain oil + gratings). **OP**: odorant phase (oil + odorant + gratings; Controls experienced 2 consecutive MOP instead). **P**: pause (darkness). Note that the training portion (the first part of the paradigm) was excluded from the analysis. Regarding the *original* versus *transformed* output from FicTrac ^[41]^ calculations: when the limit of ± 3 radians are reached, the coordinate system is flipped to the opposite sign. Inversion points were identified as those points where the first derivative exceeded 2 times the standard deviation.

Our research follows up on our previous work on the OKR and aligns with other studies on the effects of odorants on *Drosophila* flight and walking conducted in the past years ^[17–21]^. We aimed with the current work at analysing these effects within a more complete locomotion scenario - the OMR - which incorporates the proprioceptive motor feedback from the flies’ legs and needs to regulate their movement pace and patterns ^[22]^. We thus tracked the sphere’s movement to reconstruct the flies’ fictive walking paths in order to extract the trajectories and other kinematic parameters in response to the concomitant exposure of naive flies to optokinetic stimulation and to a gaseous plume carrying one of the compounds. Since it is known that aversive compounds can affect the OKR modulation in flies by decreasing the number of head-like optokinetic *nystagmi,* while leaving the intrinsic dynamics unaltered ^[23]^, we first considered whether this was also the case for the OMR. Our results show that, indeed, the compounds tested do affect the output and dynamics of the OMR, both qualitatively and quantitatively. Moreover, we found evidence that some of the synthetic masker compounds, typically found as active ingredients in some commercially available products, are not as neutral to insects as it is expected, rather they can interfere with fundamental processes underlying navigation in these (and most likely other) species.

## Pilot study’s results

### Gratings’ period, speed, and aversive odour influence the flies’ response the optomotor stimulation

First and foremost, we needed to test and tune our experimental apparatus; besides, we also wanted to set a suitable visual stimulus for the subsequent experiments. For achieving this, we conducted an initial set of experiments (Suppl. Fig. 1 and 2; Suppl. Fig 1a shows the experimental paradigm sequence) by testing gratings optomotor stimulation at different combinations of spatial period (𝜆) and velocity (𝓋), adapting the experimental paradigm found in Götz (1973) ^[24]^. In this and other studies of his, Götz analysed the visual control of locomotion on tethered *D. melanogaster* flies; in the 1973 study in particular, he had utilized the visual stimulation we replicated in order to investigate the phenomenon on flies tethered to a magnetic sledge performing stationary walking on a rubber sphere laid on servomechanisms. Two main findings of his afore cited work in particular should be mentioned here: *i*) leg contribution to the movement was observed to be equal for each pair of legs; *ii*) he identified the “limit of resolvability”, which is the minimum 𝜆 value (around 9.0° in walking flies) before observing an inversion of rotational movement (majority of movement is performed opposing the actual stimulus direction).

In our pilot set of experiments (Suppl. Fig. 1 and 2), we first tested for different sets of velocities for the gratings mask at, in our opinion, a low 𝜆 (12°) but nonetheless higher than the inversion point identified by Götz ^[15,24]^:

A. 15°.s^−1^
B. 30°.s^−1^
C. 60°.s^−1^

We observed that the slowest velocity (**A**) elicited the higher mean speed in our flies (Suppl. Fig. 1b); therefore, we kept this V value while introducing other three sets of conditions:

D. concurrent aerosol administration of a mineral oil solution containing eugenol (1% in volume) in the opposite direction of the visual stimulation
E. alternating direction of movement for the gratings (𝓋 = 15°.s^−1^, 𝜆 = 12°)
F. doubled spatial period (𝓋 = 15°.s^−1^, 𝜆 = 24°)

Although mean velocities decreased significantly in **E** and **F** (Suppl. Fig. 1b), the mean quantity of movement resulted higher than in the baseline **A** (Suppl. Fig. 1c). We measured this as ‘Rotation Index’ (**R**.**I**.), which was calculated as the measured amount of rotational movement (around the z-axis, subjects’ mean) during a trial divided by the expected amount of the same motion at set velocity in ideal conditions (*i.e.*, if the animals were moving continuously at constant speed). Mean R.I. values higher than |1| indicates that the flies turned more than expected and with a higher degree of variability in speed and acceleration than an ideal subject perfectly matching the stimulus speed and moving in a uniformly fashion. Both the **E** and the **F** condition exhibited higher index value, the latter being higher. Notably, the **D** condition gave preliminary insights on the efficacy of the substance, clearly moving mainly in the opposite direction, with respect to the **A** baseline condition, as visible in the overall rotations and velocity (Supp. Fig. 2, p < 0.0005), as well as in the mean R.I. (Suppl. Fig. 1c; p < 0.0005, see Methods).

Considering the results from the pilot experiment altogether, we selected for the next experimental paradigm the conditions which promoted the highest velocity (𝓋 = 15°.s^−1^) and amount of rotation (𝓋 = 15°.s^−1^, 𝜆 = 24°).

**Supplementary Figure 1.**
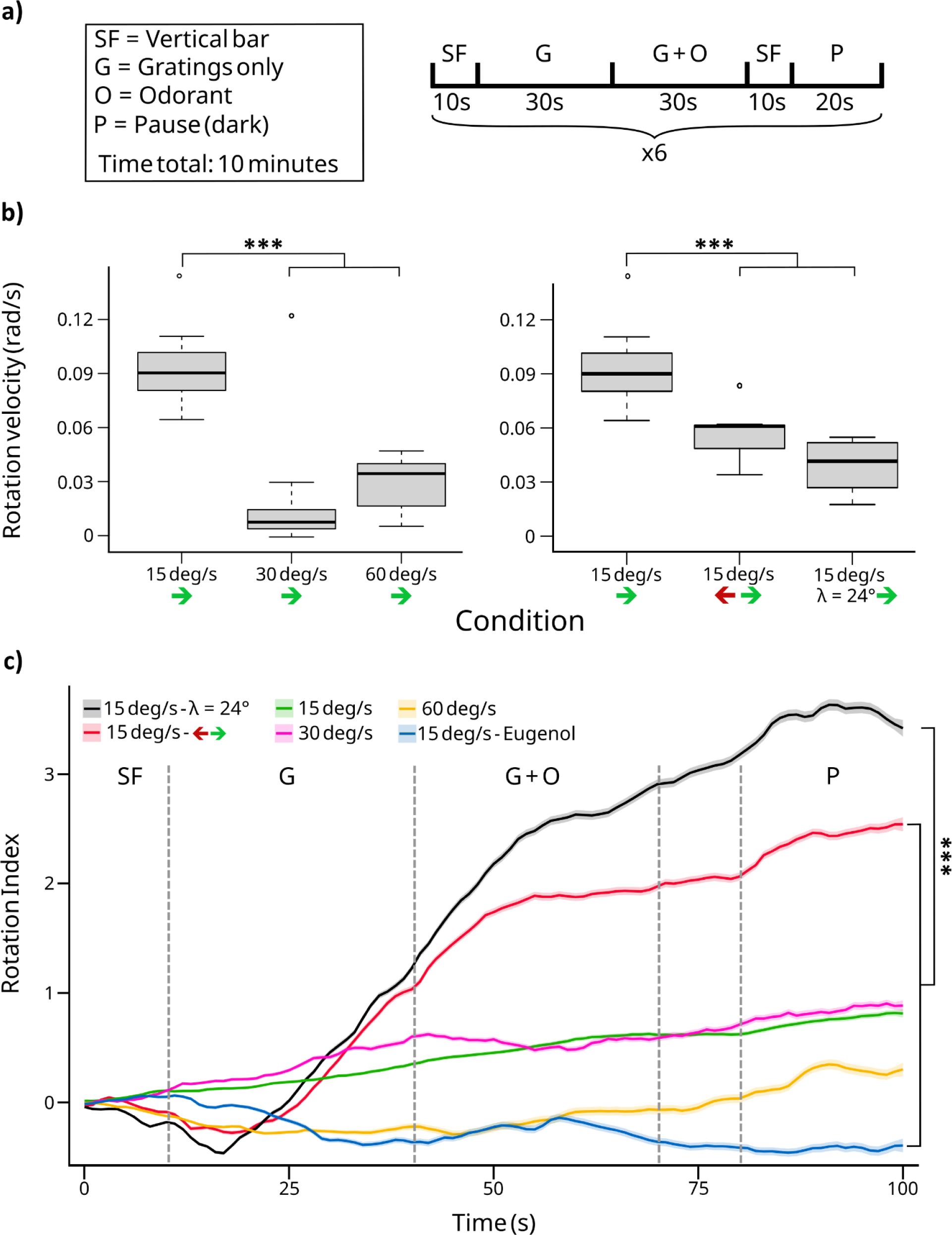
Pilot study paradigm and results. (**a**) Experimental paradigm: each trial is repeated 6 times over a span of 10 minutes and starts with a single black vertical bar (6°) on a white background for 10 seconds to centre the flies’ attention towards the middle of the screen and to elicit movement. Subsequently a mask of gratings runs for 1 minute; in the eugenol group, after 30 seconds, the air current is switched to come from a vial containing the solution with the repellent. The trial ends with 20 seconds of darkness and then a new trial starts. All conditions had 𝜆 = 12°, except for the one identified by 𝜆 = 24°. (**b**) Standard boxplots of velocity profiles. P-values: *** = 0.0005 (ANOVA + Tukey HSD test). (**c**) Rotation Index for each experimental condition (mean value among trials ± s.e.m.): the index was calculated as the mean measured amount of rotational (over the z-axis) movement during a trial divided by the theoretical amount of the same motion which would have been performed at set velocity if the animals were moving at constant speed. Values higher than |1| indicates the flies turned more than expected and with a higher degree of variability in speed and acceleration than an ideal subject perfectly matching the stimulus speed and moving in a uniformly fashion. P-values were computed on the regression line slopes: *** = 0.0005 (Tukey adjustment).

**Supplementary Figure 2.**
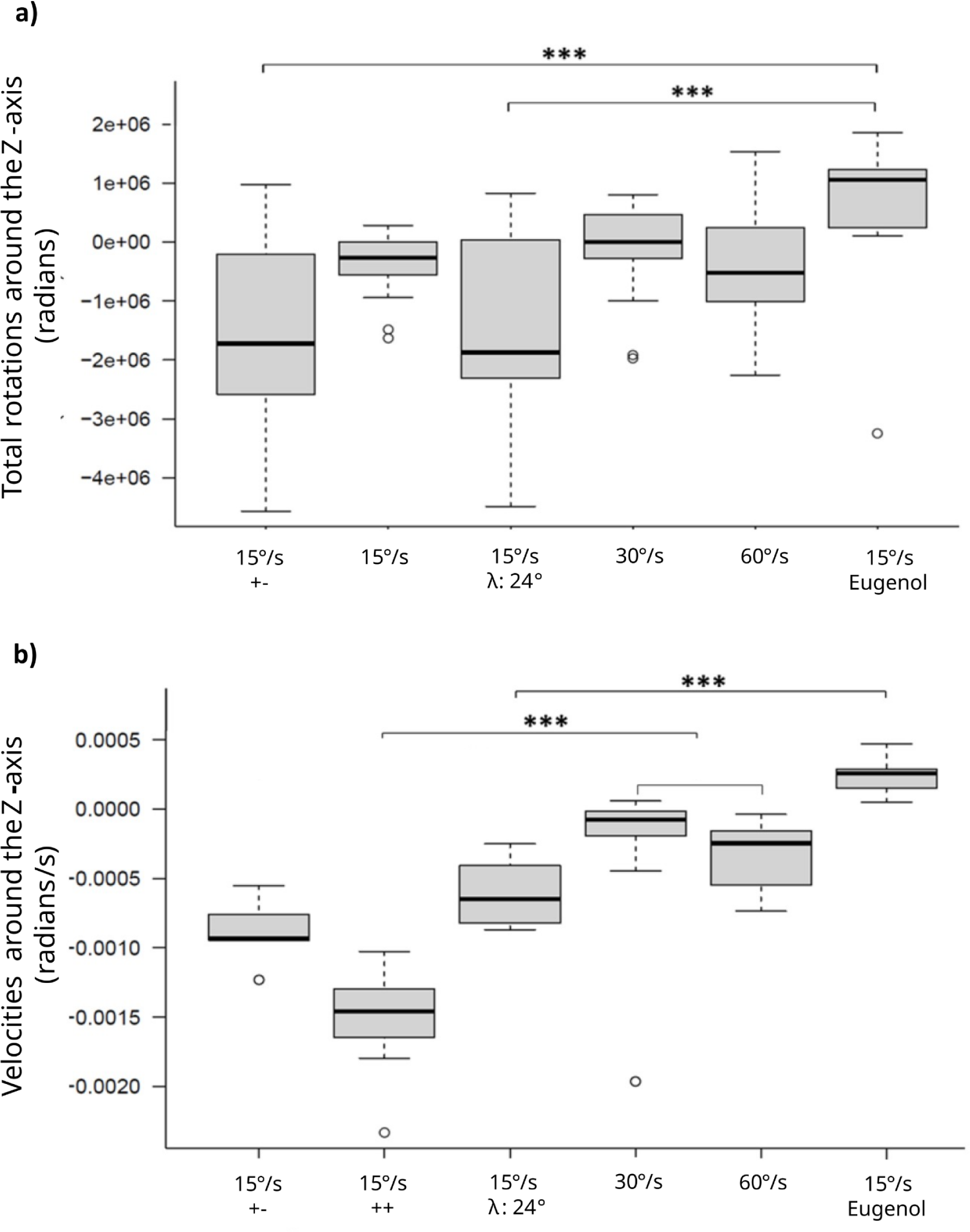
Summary of rotations and velocities. (**a**) Standard boxplots for the total amount of rotations around the vertical (Z) axis: in the “*15°·s^−1^* +-” condition the gratings move at 15°·s^−1^ in the opposite direction to the “*15°·s^−1^*” one; in the “*eugenol*” condition the gratings speed is set at *15°·s^−1^* gratings and the repellent is presented throughout the 2^nd^ half of the OK stimulation; in one condition 𝜆 was increased to 24°, instead of 12°. (**b**) Standard boxplots for the average velocity of the same conditions as in (**a**); p-values: ** = 0.005, *** = 0.0005 (ANOVA + Tukey HSD test).

## Results

### 1. Odour presentation influences the fly walking trajectories and the OMR

Initially, we aimed at assessing the relative effect of the odour presentation with respect to the concomitant visual stimulation, that, depending on the specific condition (compound and concentration), could potentially result either in a suppression or mild/strong modulation of the OMR.

We found that several groups, such as those exposed to 1% eugenol (**E1**) and 1% IR3535 (**I1**), but also 0.5% lemongrass (**L05**) and 0.5% Picaridin (**P05**), showed altered trajectories (mixed or opposite rotation direction) with respect to Controls (**C**). In particular, the analysis of the cumulative trajectories over the whole experiment (Fig. 2) and the cumulative relative number of rotations (Fig. 3) performed in the direction coherent (***+***) or non-coherent (***-***) with the grating’s direction (only taking into account the periods when the grating mask was active) revealed distinct trajectory patterns and preferred + or - direction of movement correlated with the different odorants. Suppl. Figure 3 shows the average number of (+) or (-) rotations per fly occurring in the absence (**MOP**) and in the presence (**OP**) of repellent for each group of flies.

**Figure 2.**
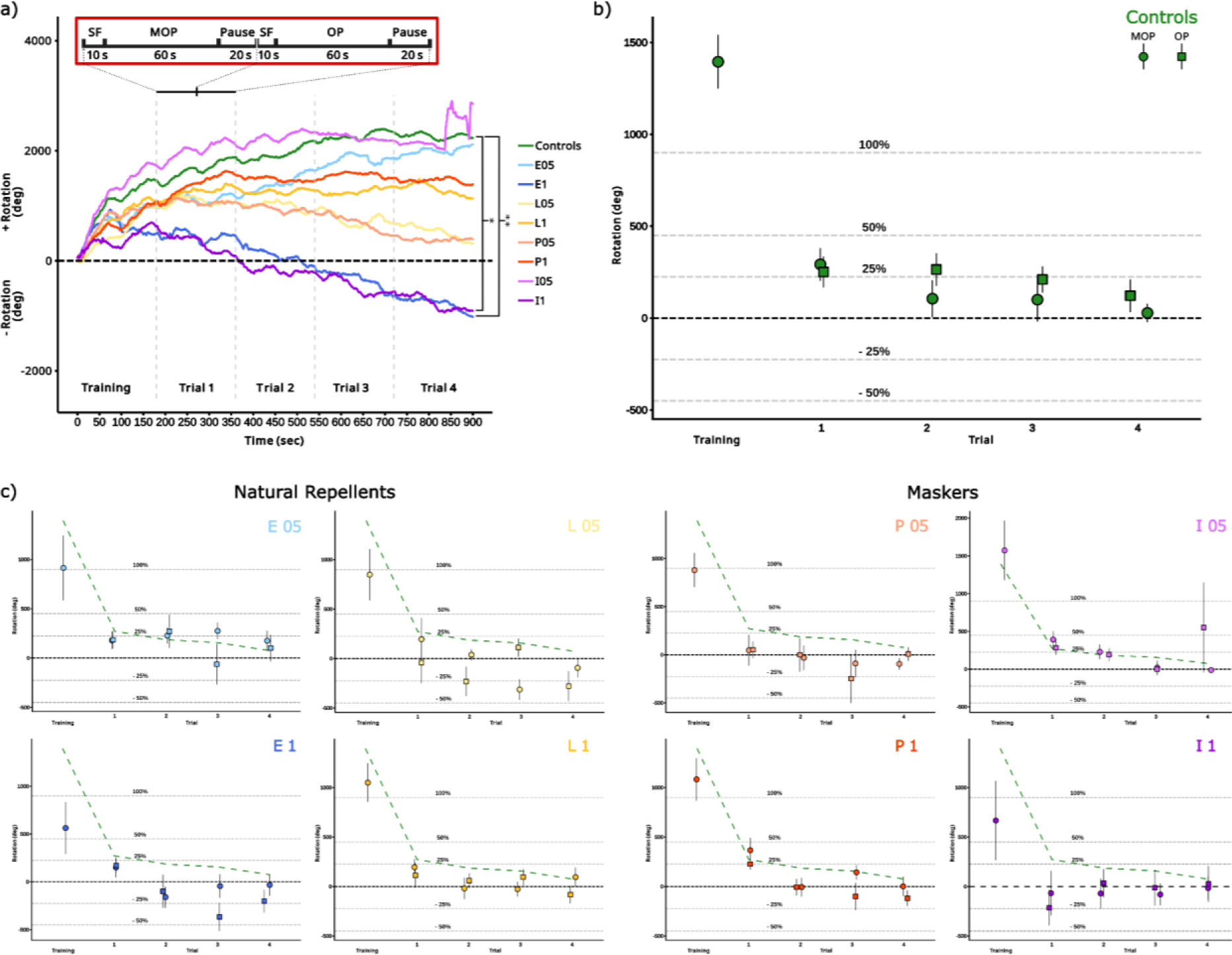
Cumulative trajectories and the average amount of rotation per trial (MOP and OP) for each compound. (**a**) Cumulative group trajectories: **L1** and **P1**, as well as **E05** and **I05** behave similarly to **C**; **P05** and **L05** have p-values close to significant (0.13 and 0.07 respectively, 0.04 and 0.02 without adjustment); **E1** and **I1** clearly tend to move in the opposite direction, away from the odour source (p-values 0.006 and 0.037 respectively). P-values were obtained from a *Kruskal-Wallis and post-hoc Dunnet test with Benjamini-Hochberg adjustment* on the subject mean differential position value each 100 milliseconds. (**b**) Average rotation per trial in the Control group. (**c**) Average rotation per trial in all the test groups; a green reference dashed line identifies the Control values shown in (**b**). * < 0.05, ** < 0.005.

**Figure 3.**
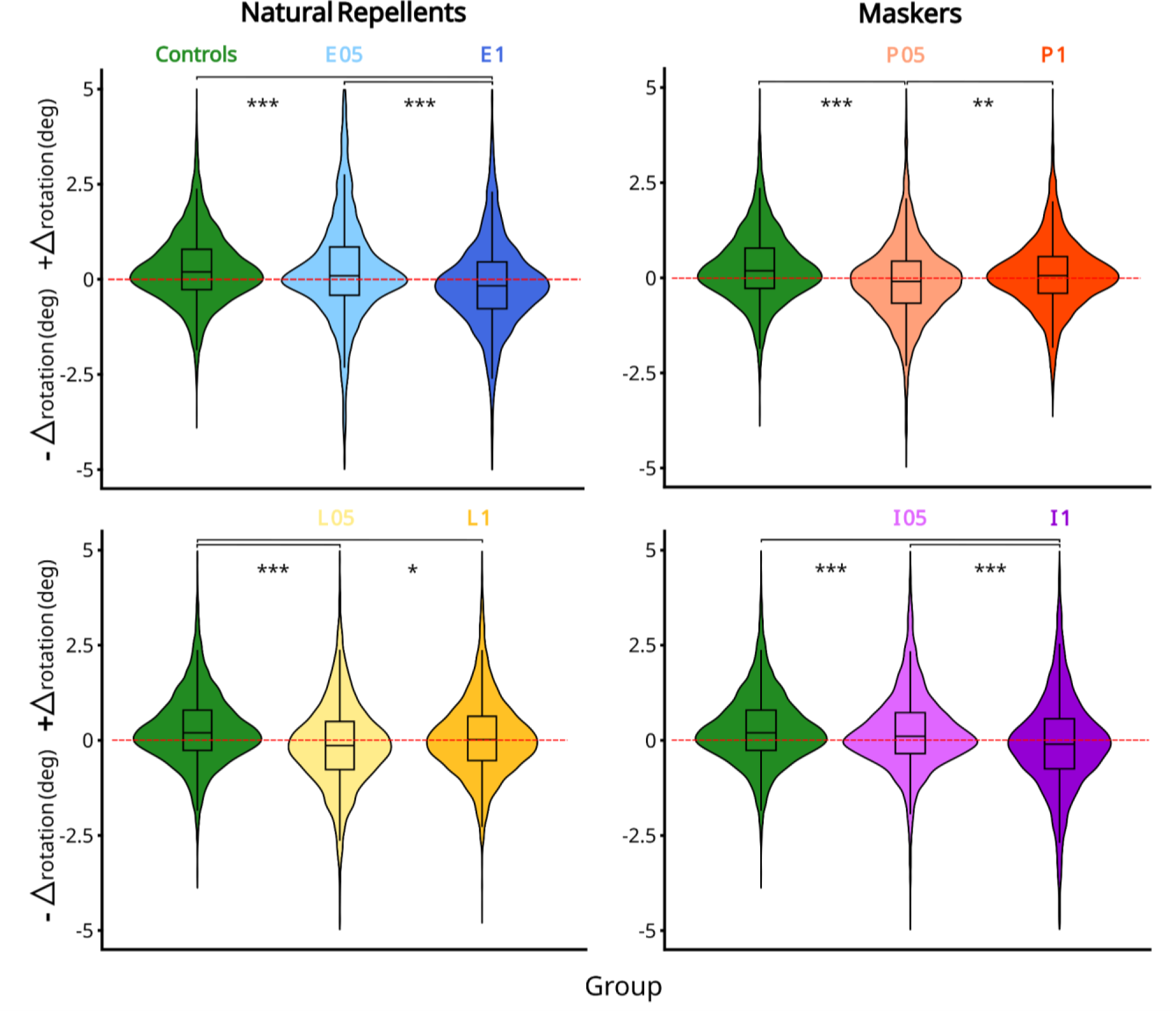
Violin plots of the Δ rotation angle. **C** flies rotate more in the direction concordant with the gratings movement, as expected. The same is true for **E05** and **I05**. **L1** and **P1**: a slight preference towards ***concordant*** (***+***) directions can be observed (see Suppl. Fig. 3 for the MOP/OP division). **E1**, **L05**, **P05** and **I1** all show a tendency for the ***opposite*** (***–***) directions both during MOP and OP, although this is visible as ***opposite*** (***–***) overall <u>trajectories</u> only in **E1** and **I1** (Fig. 2a). *: p < 0.05, ***: p < 0.0005: see **Table 1** for detailed number of rotations and p-values (pairwise comparison of proportion with Bonferroni p-value adjustment).

**Supplementary Figure 3.**
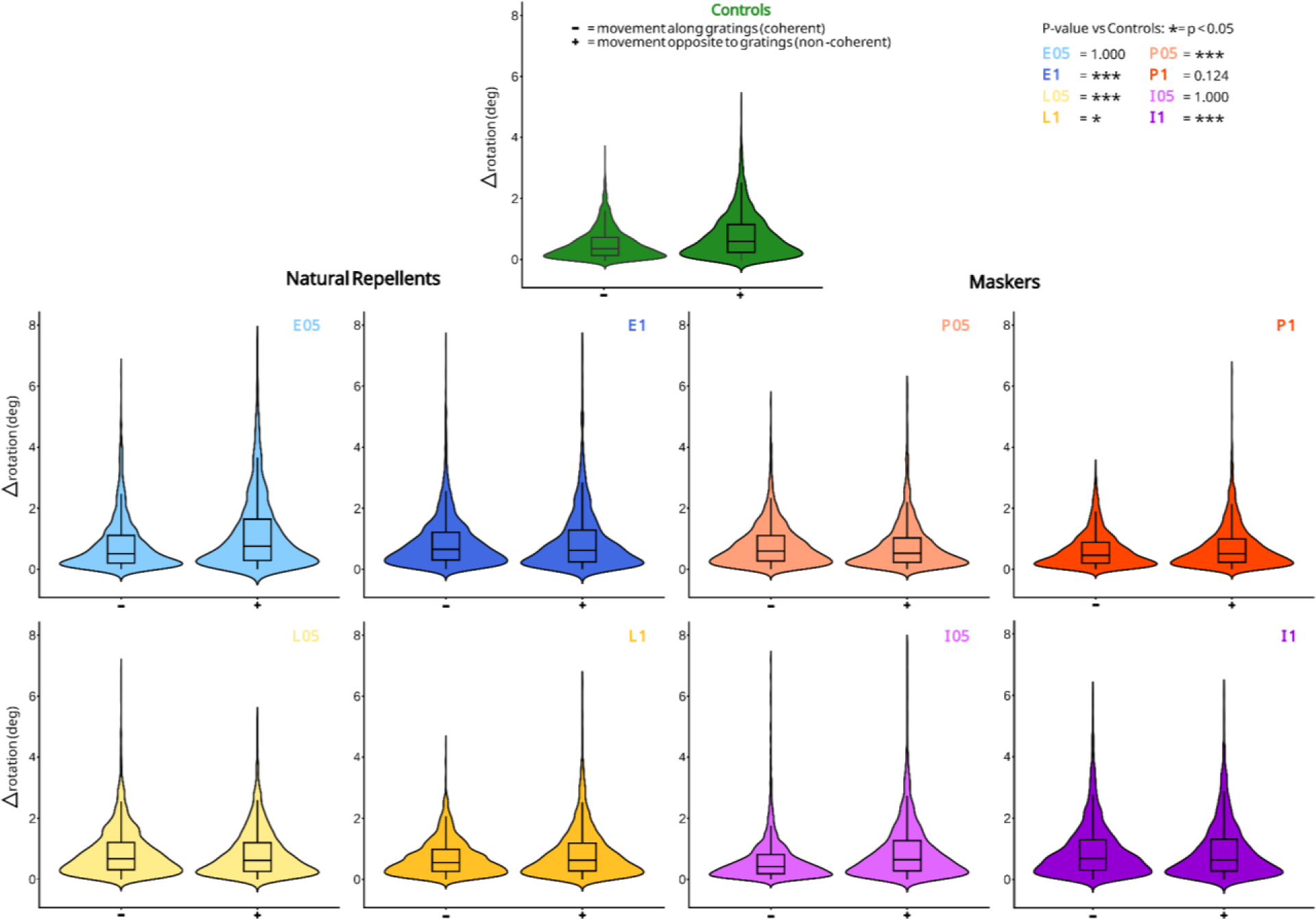
Violin plots of the Δ rotation angle during MOP and OP phases. Same plot as in Figure 3, except MOP and OP are shown separately.

**Table 1.** Concordant vs discordant number of rotations. Values relative **Fig. 3** panels. Cumulative values (mean ± standard deviation calculated on the variation in position every 100 milliseconds) per group: trajectories include the training period (first 180 seconds). Values for the mean amount of movement concordant (***+***) or discordant (***-***) with the gratings’ direction only include the proper trials (1 ∼ 4) during which the grating mask is projected on screen. P-values related to the relative +/− rotation proportions are obtained through pairwise comparison of proportions with Bonferroni correction.

| Group | Trajectory<br>(mean rotations) | coherent<br>movement<br>(mean + rotations) | non-coherent<br>movement<br>(mean - rotations) | P-value<br>vs<br>Controls |
| --- | --- | --- | --- | --- |
| Controls (n = 13) | 6. ± 2.1e-3 | 6.71 ± 2 e-3 | 2.87 ± 1.4e-3 | - |
| Eugenol 0.5% (n = 7) | 5.72 ± 3.1e-3 | 8.56 ± 3.3e-3 | 4.80 ± 2.4e-3 | 1.000 |
| Eugenol 1% (n = 10) | -2.98 ± 2.8e-3 | 5.41 ± 2.6e-3 | 6.78 ± 2.3e-3 | $< 2e^{-16}$ |
| Lemongrass 0.5% (n = 10) | 0.72 ± 2.3e-3 | 4.89 ± 2.1e-3 | 6.59 ± 2.1e-3 | 6.3e<sup>-10</sup> |
| Lemongrass 1% (n = 12) | 2.98 ± 2.1e-3 | 5.75 ± 2.2e-3 | 4.57 ± 1.6e-3 | 0.008 |
| Picaridin 0.5% (n = 11) | 0.92 ± 2.2e-3 | 4.65 ± 2.1e-3 | 5.63 ± 1.9e-3 | 1.7e<sup>-11</sup> |
| Picaridin 1% (n = 12) | 3.69 ± 1.9e-3 | 5.23 ± 1.9e-3 | 3.83 ± 1.6e-3 | 0.124 |
| IR3535 0.5% (n = 10) | 7.77 ± 6.1e-3 | 8.82 ± 6.9e-3 | 4.22 ± 3. 6e-3 | 1.000 |
| IR3535 1% (n = 13) | -2.69 ± 2.6e-3 | 5.64 ± 2.4e-3 | 6.78 ± 2.4e-3 | $< 2e^{-16}$ |

Considering the mean cumulative trajectories during the whole task (Fig. 2a), the repellent **E1** (n = 10, p-value = 0.006) and the masker **I1** (n = 10, p-value = 0.037) showed major differences from the **C** (n = 13) group, exhibiting the opposite rotation direction, which is non-coherent with the experienced visual stimulation and moving away from the odour source. In contrast, **E05** (n = 7), **L1** (n = 12), **P1** (n = 12), and **I05** (n = 13) are not significantly different with respect to **C**. Moreover, **L05** (n = 10) and **P05** (n = 11), approached statistically significant differences with respect to **C** (p-values = 0.07 and 0.13 respectively), showing an intermediate behaviour with respect to the controls and the groups showing the most non-coherent rotations (*Kruskal-Wallis: chi-squared = 26.104, d.f. = 8, p-value = 0.001. Post-hoc: Dunnet test with Benjamini-Hochberg correction*).

In particular:

- The repellent Eugenol and the masker IR3535 show a similar trend of concentration-dependent impact: at low concentration it does not appear on the walked trajectories any clear repellent effect, rather an opposite trend if any. However, the repellent action becomes detectable and significant at higher concentrations.
- The repellent Lemongrass and the masker Picaridin show an opposite trend, characterized by a significant repellent effect at lower concentrations, while showing a diminished if not opposite effect at higher concentrations, possibly due to saturation.

These differences in the cumulative trajectories were confirmed by the cumulative mean amount of movement observed in the direction coherent (+) or non-coherent (-) with the visual stimulation (Fig. 3). Here, without considering the initial 180 seconds of the training period, both **E1** and **I1** exhibited a reversed ratio for the preferred direction of movement with respect to **C**. Furthermore, a similar, significant, difference was also observed for **L05** and **P05** (see Table 1 for the mean values ± standard deviations alongside the Bonferroni adjusted p-values for the equality of proportions test).

When considering the mean values by subject (Suppl. Table 1), the trends observed for the cumulative trajectories and relative quantity of (+/−) rotations were confirmed. No significant differences were found between MOP and OP for any group: **C** flies moved more in the direction coherent with the gratings, as expected, as well as **E05** and **I05**. Moreover, while **L1** and **P1** had a slight preference towards (+) directions during the MOP – which reversed during OP – **E1**, **L05**, **P1** and **I1** all preferred the (-) direction (*Kruskal-Wallis: chi-squared = 4.7795, d.f. = 17, p-value = 0.9983*), while significant differences were present for the preferred direction of rotation (*Kruskal-Wallis: chi-squared = 306.25, d.f. = 35, p-value < 2.2e^−16^. Post-hoc: Dunnet test with Benjamini-Hochberg correction*).

**Supplementary Table 1.** Average per fly concordant vs discordant number of rotations. Values shown as mean ± sd. In all groups MOP and OP do not present statistically significant differences (*Kruskal-Wallis: chi-squared = 4.7795, d.f. = 17, p-value = 0.9983*). Conversely, direction within MOP or OP is significant (*Kruskal-Wallis: chi-squared = 306.25, d.f. = 35, p-value < 2.2e-16 + Dunnet test with Benjamini-Hochberg correction*).

| Group | MOP |  |  | OP |  |  |
| --- | --- | --- | --- | --- | --- | --- |
|  | Mean<br>- rotations | Mean<br>+ rotations | P-value | Mean<br>- rotations | Mean<br>+ rotations | P-value |
| Controls (n = 13) | 10.06 ± 1.81 | 12.71 ± 3.93 | 0.00140 | 10.64 ± 2.54 | 12.54 ± 3.96 | 0.00096 |
| Eugenol 0.5% (n = 7) | 12.85 ± 1.51 | 15.68 ± 4.23 | 0.00048 | 12.60 ± 2.33 | 15.56 ± 4.61 | 0.00019 |
| Eugenol 1% (n = 10) | 15.10 ± 4.73 | 12.05 ± 1.46 | 0.00007 | 14.51 ± 2.29 | 11.07 ± 1.53 | 0.00077 |
| Lemongrass 0.5% (n = 10) | 15. 21 ± 4.14 | 14.06 ± 4.55 | 0.00045 | 15.28 ± 4.20 | 13.06 ± 2.97 | 0.00004 |
| Lemongrass 1% (n = 12) | 12.63 ± 4.28 | 13.63 ± 4.28 | 0.00006 | 12.76 ± 2.85 | 12.48 ± 2.70 | 0.00007 |
| Picaridin 0.5% (n = 11) | 13.69 ± 5.64 | 11.80 ± 2.50 | 0.00028 | 14.33 ± 5.46 | 12.55 ± 3.60 | 0.00009 |
| Picaridin 1% (n = 12) | 10.58 ± 3.32 | 11.89 ± 3.25 | 0.00250 | 10.88 ± 3.65 | 10. 85 ± 2.09 | 0.00520 |
| IR3535 0.5% (n = 10) | 11.46 ± 2.00 | 12.74 ± 2.79 | 0.00009 | 14.27 ± 8.58 | 16.11 ± 15.11 | 0.00006 |
| IR3535 1% (n = 13) | 15.48 ± 8.43 | 12.28 ± 3.31 | 0.00015 | 15.28 ± 7.93 | 11.81 ± 3.41 | 0.00025 |

### 2. Mean rotation velocity increases as flies move against the direction of gratings

Next, we evaluated whether the presence of repellents could influence the dynamics of the rotational locomotor pattern induced by the visual stimulation and analysed the velocity profiles for each group of flies. First, for each group, we computed the average angular velocity 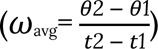 with respect to the z-axis during the presentation of visual stimulus (moving gratings on screen) across the 4 trials (Fig. 4 and Tables 2 – 3). Decomposing the data according to the movement directionality, revealed that, for the *coherent* (**+**) direction, with respect to the control condition, only **E05** group shows a higher angular velocity. For the *coherent* (**+**) direction this effect was, however, not detected when further segregating the data according to the phase (MOP or OP), possibly because of the relatively smaller sample size. The velocity profiles of all the other groups were similar (*3W-Robust ANOVA: “group”, F = 17.04, p-value = 0.061; “direction”, F = 3969.60, p-value = 0.0001; “group: direction”, F = 34.93, p-value 0.0010; “stim (MOP/OP)” and related interactions not significant. Post-hoc: mcppb20 test with Bonferroni correction*).

**Figure 4.**
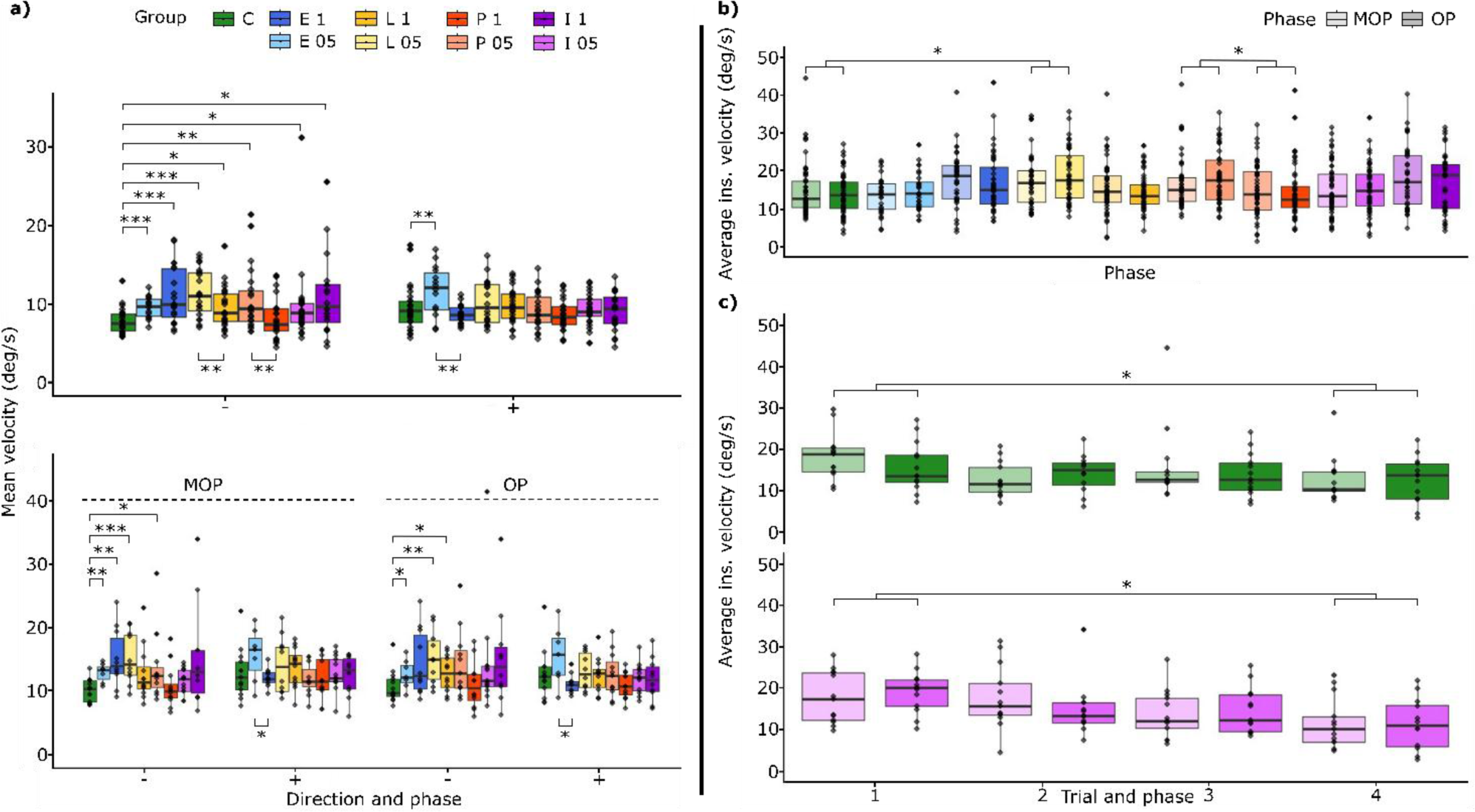
Mean velocity and averaged instantaneous velocities. **Legend**: group colours are shown in the upper left panel with the same label in the main text; in panels **b, c, d,** degree of transparency relates to MOP (lighter) and OP (darker). (**a**) **Top left**: standard boxplots for the mean speed of flies during the optomotor stimulation in both directions, concordant (+) or discordant (-) with the gratings’ movement. **Bottom left**: same plot separated between MOP and OP (test: *2W* or *3W-Robust ANOVA + post-hoc mcppb20 test with Bonferroni correction*, see main text and **Tables 2** – **3**). (**b**) Standard boxplots for the averaged instantaneous velocity mediated by subject (see Table 4); (**c**)Average ins. velocity along trials of Controls and IR3535 0.5% respectively (test: *2W-ANOVA for Aligned Rank Transformed Data + EMMs pairwise with Bonferroni correction*, see main text). P-values: * < 0.05, ** < 0.005, *** < 0.0005.

**Table 2.** Average MOP and OP rotation velocities. Values relative to **Fig. 4a** shown as mean ± SD. P-value(s) obtained from within-group paired t-tests on MOP/OP data (Eugenol 1%: t = −2.58 - d.f. = 9).

| Group | Mean rotation velocity (deg-sec-1) |  |  |  |  |  |
| --- | --- | --- | --- | --- | --- | --- |
|  | MOP |  |  | OP |  |  |
|  | - | + | P-value | - | + | P-value |
| Controls (n = 13) | 7.55 ± 0.22 | 9.53 ± 0.47 | ns | 7.98 ± 0.30 | 9.41 ± 0.47 | ns |
| Eugenol 0.5% (n = 7) | 9.64 ± 0.18 | 11.76 ± 0.50 | ns | 9.45 ± 0.28 | 11.67 ± 0.55 | ns |
| Eugenol 1% (n = 10) | 11.33 ± 0.56 | 9.04 ± 0.17 | ns | 10.88 ± 0.63 | 8.30 ± 0.18 | 0.0296 |
| Lemongrass 0.5% (n = 10) | 11.40 ± 0.49 | 10.55 ± 0.54 | ns | 11.46 ± 0.50 | 9.79 ± 0.35 | ns |
| Lemongrass 1% (n = 12) | 9.47 ± 0.51 | 10.24 ± 0.37 | ns | 9.57 ± 0.34 | 9.36 ± 0.32 | ns |
| Picaridin 0.5% (n = 11) | 10.27 ± 0.67 | 8.85 ± 0.30 | ns | 10.75 ± 0.65 | 9.41 ± 0.43 | ns |
| Picaridin 1% (n = 12) | 7.94 ± 0.40 | 8.92 ± 0.39 | ns | 8.16 ± 0.43 | 8.14 ± 0.25 | ns |
| IR3535® 0.5% (n = 10) | 9.45 ± 0.24 | 9.56 ± 0.33 | ns | 10.70 ± 1.02 | 12.08 ± 1.80 | ns |
| IR3535® 1% (n = 13) | 11.61 ± 1.00 | 9.21 ± 0.39 | ns | 11.46 ± 0.95 | 8.86 ± 0.41 | ns |

**Table 3.**
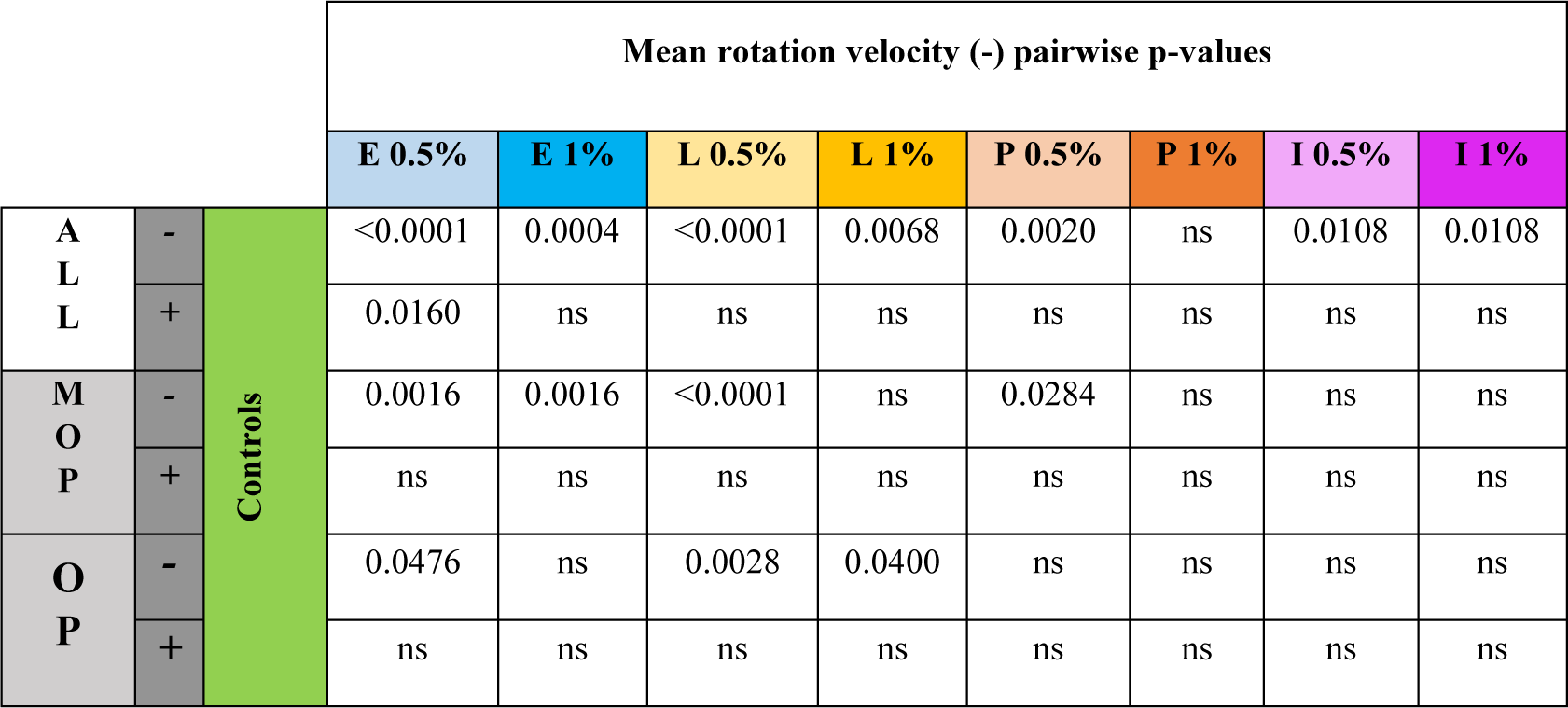
P-values from the Robust ANOVA and the post-hoc mcppb20 tests. Significance relative to **Fig. 4a**. To reduce the effect of outliers the tests (*Robust ANOVA*, see text and Methods) were run after trimming the outliers with values present in the bottom (<10%) and top (>90%) of the data. Post-hoc mcbbp20 tests p-values were adjusted with Bonferroni correction for multiple comparisons.

|  |  |  | Mean rotation velocity (-) pairwise p-values |  |  |  |  |  |  |  |
| --- | --- | --- | --- | --- | --- | --- | --- | --- | --- | --- |
|  |  |  | E 0.5% | E 1% | L 0.5% | L 1% | P 0.5% | P 1% | I 0.5% | I 1% |
| A<br>L<br>L | - | Controls | <0.0001 | 0.0004 | <0.0001 | 0.0068 | 0.0020 | ns | 0.0108 | 0.0108 |
|  | + |  | 0.0160 | ns | ns | ns | ns | ns | ns | ns |
| M<br>O<br>P | - |  | 0.0016 | 0.0016 | <0.0001 | ns | 0.0284 | ns | ns | ns |
|  | + |  | ns | ns | ns | ns | ns | ns | ns | ns |
| O<br>P | - |  | 0.0476 | ns | 0.0028 | 0.0400 | ns | ns | ns | ns |
|  | + |  | ns | ns | ns | ns | ns | ns | ns | ns |

**Table 4.**
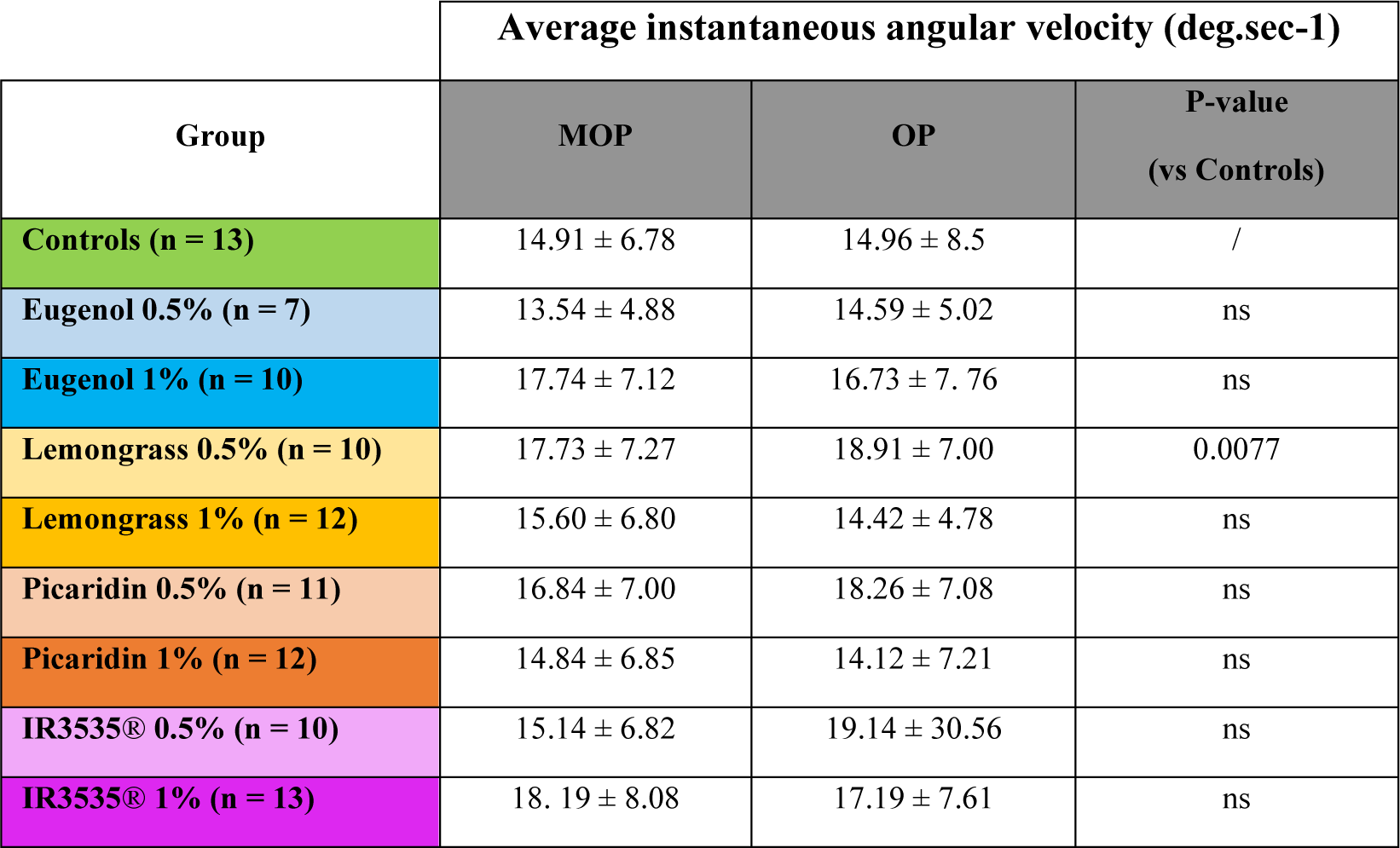
Mean and standard deviation values for the instantaneous angular velocity. Values referring to **Fig. 4b**. No significant differences were found between **MOP** and **OP**. The only significant difference with respect to **C** is observed for **L05**; **P05** and **P1** are the only pair with a within group significant difference (p = 0.0476). P-values extracted from *Aligned Rank Transformation ANOVA + post-hoc Estimated Marginal Means with Pairwise Comparisons (EMMs) and Bonferroni correction*, see main text.

In contrast, for the *non-coherent* (**-**) direction, several differences could be appreciated: apart from **P1**, all groups showed increased velocity of rotation with respect to controls. Moreover, while **E05** – **E1** and **I05** – **I1** didn’t show a significant difference between concentrations (**E1** and **I1** values were more spread), this was instead true for **L05** – **L1** and **P05** – **P1**. When separating between MOP and OP, we found that **E05** and **L05** exhibit increased velocity during both phases, while in the case of **P05** and **E1** the major contribution came from the MOP. **L1**, conversely, moved faster only when the repellent was actively delivered to flies during the OP.

### 3. Overall instantaneous velocity profiles remain unchanged, with a trend towards reduced values as trials proceed

Next, we wondered whether during the protocol flies could show effects due to muscular fatigue. We then calculated the instant angular velocities 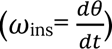 integrating the information from all the three axes of rotation (Table 4). After testing, the overall 𝜔_ins_ profiles were similar across all groups, except for **L05** (p-value = 0.0077), which showed higher speeds with respect to **C** (see Fig. 4b and Table 4). For thoroughness, on inspection of the plot **P05** and **I1** also appeared to stand out, but the p-values were just short of significance (0.0733 and 0.0749 respectively). Picaridin was the only repellent showing a significant difference between the two tested concentrations, with **P05**, the lower one, showing higher instantaneous velocity (p-value = 0.0476, *3W-ANOVA for Aligned Rank Transformed Data: “group”, F = 4.59, p-value = 1.75e^−5^; “trial”, F = 3.68, p-value = 0.0118; “stim (MOP/OP)” and interactions not significant. Post-hoc: EMMs pairwise comparison with Bonferroni correction*).

Additionally, as the statistical test showed a slightly positive dependence from the “trial” variable, we analysed each group and found a positive correlation in the **C** and **I05** group. For both (see Fig. 4c – 4d), the velocity values showed a slight progressive reduction along the trial sequence, with a significant difference between the 1st and the 4th trial for both **C** (p-value = 0.0325) and **I05** (p-value = 0.0011) which also revealed a positive response between the 1st and 3rd trial (p-value = 0.0407). No other group exhibited a similar trend (*2W-ANOVA for Aligned Rank Transformed Data: “trial”, F = 3.49, p-value = 0.0185; “stim” (MOP/OP) and interaction not significant + EMMs pair-wise comparison with Bonferroni correction*) and **C** group(*2W-ANOVA for Aligned Rank Transformed Data: “trial”, F = 3.49, p-value = 0.0426; “stim” (MOP/OP) and interaction not significant*).

### 4. Low concentrations of natural repellents reduce the time interval between bursts of movement

Following the observations on the average velocity, we reasoned that the higher velocity found during Eugenol and Lemongrass (all concentrations) stimulation might be due to an increased sensory engagement causing livelier patterns of movement. We thus checked whether repellent delivery was affecting how frequently flies alternated pauses and bouts of locomotor activity, and how long these periods of activity/inactivity lasted.

For this purpose, we calculated the angular acceleration values 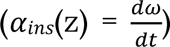 of the trajectory around the z-axis to extract “*go*” and “*stop”* epochs based on the relative acceleration values with respect to the time averaged value. We found that, across the 4 trials, the average number of “*stop*” and “*go*” did not change between groups (see Fig. 5b, further down); also, the average duration of the movement bouts (Fig. 5a, top) did not show appreciable differences with respect to **C** (MOP: mean 369.37 ± SD 56.20 milliseconds, OP: 370.40 ± 48.37 ms) except in the case of the **P05** group (MOP: 336.47 ± 40.67 ms, OP: 334.12 ± 49.86 ms), which exhibited slightly shorter bouts (p-value = 0.0003, *2W-ANOVA for Aligned Rank Transformed Data: “group”, F = 6.89, p-value = 8.79e-9; “stim (MOP/OP)” and interaction not significant+ EMMs pairwise comparison with Bonferroni correction*). On the other hand (Fig. 5a, bottom), the average duration of the “*stop*” periods was reduced for low concentrations of the natural repellents **E05** and **L05** (MOP: 171.15 ± 30.89 ms, OP: 174.80 ± 30.29 ms; p-value = 0.0042; MOP: 180.43 ± 55.94 ms, OP: 183.85 ± 42.51 ms; p-value = 0.0266) with respect to **C** (MOP: 199.17 ± 38.90 ms, OP: 196.51 ± 42.58 ms) but without appreciable difference between the MOP and OP (*2W-ANOVA for Aligned Rank Transformed Data: “group”, F = 4.24, p-value = 5.41e^−5^; “stim (MOP/OP)” and interaction not significant + EMMs pairwise comparison with Bonferroni correction*).

**Figure 5.**
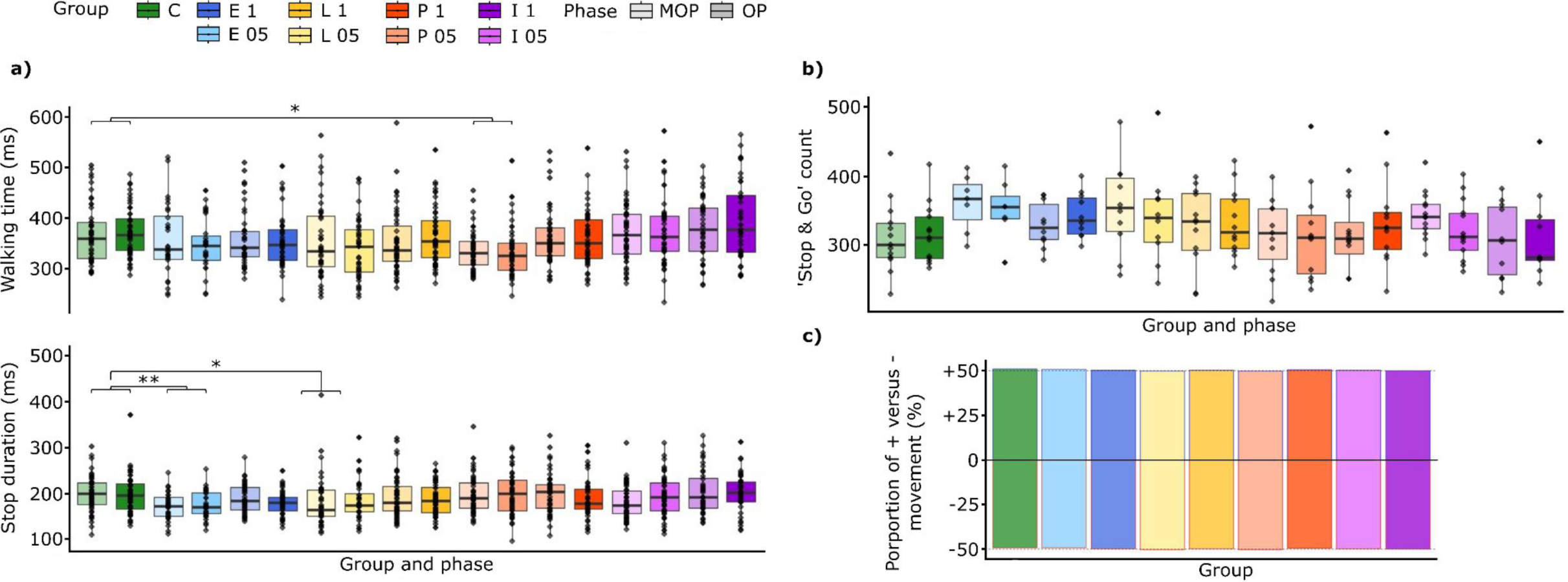
Mean velocity and averaged instantaneous velocities. **Legend**: group colours are shown in the upper left panel with the same label in the main text; in panels **a** and **b,** the degree of transparency relates to MOP (lighter) and OP (darker). (**a**) Standard boxplots for the mean duration (milliseconds) of walking (“Go”) and idle (“Stop”) periods (tests: *2W-ANOVA for Aligned Rank Transformed Data. (**a**): “group”, F = 6.89, p-value = 8.79e-9; “stim (MOP/OP)” and interaction not significant + EMMs pairwise comparison with Bonferroni correction*; *(**b**) “group”, F = 4.24, p-value = 5.41e-5; “stim (MOP/OP)” and interaction not significant+ EMMs pairwise comparison with Bonferroni correction [see main text for significant post-hoc p-values]*). (**b**) Standard boxplots for the “Go” and “Stop” count, mediated between subjects, in each group and phase (test: *Pearson’s Chi-squared X = 2.07 - p = 0.97*). (**c**) Proportion of the direction of the first movement at the onset of walking episodes concordant (***+***) or discordant (***-***) with the gratings’ direction of movement. (test: *Pearson’s Chi-squared X = 5.16 - p = 0.74* respectively). P-values: * < 0.05, ** < 0.005.

### 5. The direction of the acceleration at movement onset is random and not influenced by odour presentation

One last parameter we considered was the orientation of each first bout of movement performed by the flies after a “stop” while the visual stimulation was present (Fig. 5c). Our goal was to see whether initiation of movement could be directed and shaped by the interaction with the concomitant odour. In this respect our expectation was to see a decisive shift away from the odour, at least in those groups which were showing a decisively negative (**E1**, **I1**) or at least mixed (**L05**, **L1**, **P05**, **P1**) response with respect to the repellent.

In the end, we did not find any significant differences: in each group, the movement onset direction seemed to initiate randomly at the beginning of locomotion bouts.

## Discussion

A resume of the results we observed in our experiment is shown in Table 5 below.

**Table 5.**
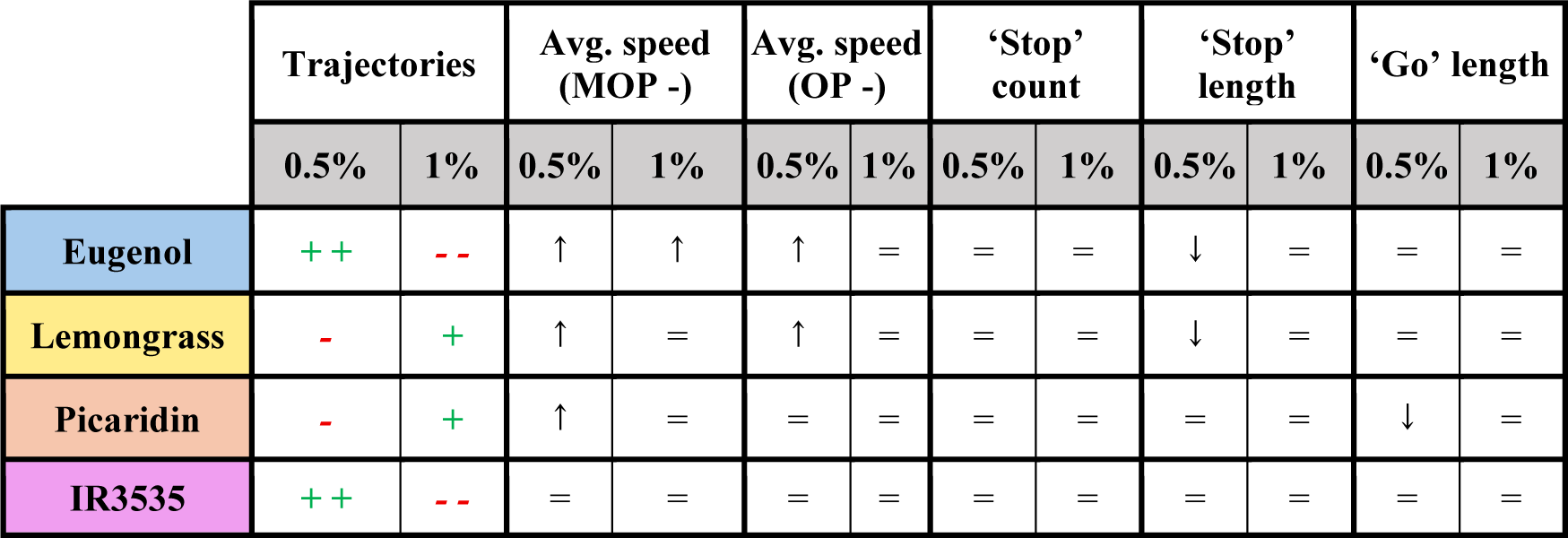
Recap of the results for the walking experiment. Red “-” indicates the direction discordant with the grating mask movement, while the green “+” is for concordant directions; “=” means no change is observed.

|  | Trajectories |  | Avg. speed (MOP -) |  | Avg. speed (OP -) |  | 'Stop' count |  | 'Stop' length |  | 'Go' length |  |
| --- | --- | --- | --- | --- | --- | --- | --- | --- | --- | --- | --- | --- |
|  | 0.5% | 1% | 0.5% | 1% | 0.5% | 1% | 0.5% | 1% | 0.5% | 1% | 0.5% | 1% |
| Eugenol | ++ | -- | ↑ | ↑ | ↑ | = | = | = | ↓ | = | = | = |
| Lemongrass | - | + | ↑ | = | ↑ | = | = | = | ↓ | = | = | = |
| Picaridin | - | + | ↑ | = | = | = | = | = | = | = | ↓ | = |
| IR3535 | ++ | -- | = | = | = | = | = | = | = | = | = | = |

From the qualitative point of view, it stands out immediately that the presence of repellents alters the prevalent direction of movement of flies undergoing the OMR stimulation. As mentioned in the Methods section, we decided *a priori* to exclude from the analysis those animals which did not show a pattern of rotation consistent with the OMR during the training phase and, therefore, it is safe to assume that the observed magnitude of *contra*-gratings movement was indeed elicited by the aversive effect of the delivered odours.

Most of the differences are found within the natural repellent groups (Eugenol and Lemongrass), but at least some of the measured parameters are indicative of an unexpected behaviour in response to the so-called maskers. Regarding the natural repellents, flies exposed to highest (1%) Eugenol concentration show a marked alteration in their walked path, moving away from the repellent source (in the opposite direction of the OMR stimulus). A similar effect, altough not significative is observed in the lowest (0.5%) Lemongrass concentration. As per the maskers, picaridin-exposed flies show mixed trajectories and suggest a possible perturbation in their action planning – the consequent motor programs – similarly to the flies experiencing the proper repellents. On the other hand, many of the analysed features do not appear as altered in flies exposed to IR3535: notably, while low concentrations of this masker do not alter trajectories with respect of the control group, the higher concentration (1%) stands out showing a strongly opposite direction of rotation with respect to Controls, just as in the case of the highest eugenol concentration.

Another noteworthy aspect is the difference shown by the two concentrations utilized for all odorants in our experiments. When such concentration-related mixed responses are detected, a trend appears for a stronger/relevant response at the lower concentrations tested. Again, we tested compounds (Eugenol, Lemongrass, Picaridin, and IR3535) and concentrations (0.5% and 1%), which proved to be functional for studying the response to the OKN stimulation. In turn, the results on the OMR here suggest that the modulation acting on the final behavioural output is possibly not peripheral, but could be highly sensitive to low concentrations, thus resulting in the rapid saturation of the system. However, another possible, and not necessarily exclusive, explanation may be that what we are observing is not a “simple” variation in the quantitative aspect of a single response, but rather that different outputs are selected among a possible repertoire of behaviours, depending on the perceived strength of the odorant. The conceivable uncertainty present in conditions of attenuated olfactory stimulation – which would hypothetically involve a greater level of integration and a consequently larger “computational” load in the selection of the most suitable behaviour – would be lost at system saturation, promoting a more decisive and unequivocal response, much as we observed in the comparison between the low and high eugenol concentrations.

In fact, we did not find any difference between the MOP and OP phases within each combination of odorant and concentration, as instead was the case in our previously reported OKR-flight paradigm ^[23]^. This is likely due to the greater complexity of the control mechanisms of the OMR, which rather than being immediately impacted by the administration of the odorants is possibly remodelled in a slower and perhaps more constrained manner ^[25]^.

In our OMR task, flies can be seen as engaged in a response guided towards a preferred direction by the elicited optic flow. Something similar, although not OMR related, was described in Frighetto *et al*. (2019) ^[26]^: the presentation of a (visual) distractor was sufficient to cause a temporary disengagement from a Buridan paradigm. In our results it may be that this is not happening during the OMR (as was instead previously observed by us to occur in the case of the OKR ^[23]^) in which case the response was not suppressed but rather modulated. The odours act as distractors involving a different sensory pathway, which may be possibly harder to integrate than a second visual distractor. The more complex responses appreciated during the OMR could be therefore explained by a remodulation or reshaping occurring within an already engaged motor program, instead of a process involving the selection of a new action after disengaging from the ongoing one.

Indeed, in these experiments, OMR is observed during grounded locomotion, which is the complex result of the interaction between the local thoracic circuits, called CPGs, generating spatio-temporally organised patterns of movements of the animal appendages and their modulation by both the cerebral ganglia, the central complex (CX), the mushroom bodies (MB) and the gnathal ganglia, as reviewed in Emanuel *et al*. (2020) ^[27]^.

It is known that the cerebral ganglia, which are centres integrating both visual and olfactory information, are involved in the regulation of probably mostly inhibitory walking behaviours, as demonstrated in experiments where insects from different species deprived of the circumesophageal connectives – which link the gnathal and cerebral ganglia – have long and unoriented bouts of movement ^[27]^. In fruit flies, the inhibitory effect of the MB was demonstrated by Martin, Ernst and Heisenberg (1998) ^[28]^, who observed highly increased walking activity in flies with ablated MB or defective Kenyon cells; meanwhile, Serway *et al*. (2009) ^[29]^ and Mabuchi *et al*. (2016) ^[30]^ demonstrated the MB role in enhancing activity when initiating light-evoked walking and maintaining the physiological daily rhythmic activity ^[27,29,30]^.

Concurrently, it has been shown that mutant flies with alterations of the CX’s subunits present reduced walking activity in terms of unreliability of movement initiation and reduced speed or increased lag between consecutive bouts ^[27,31–33]^. Moreover, in cockroaches, it has been demonstrated that a reduction in the CX’s synaptic drive diminishes the excitatory activity on the thoracic CPGs with decreased activity of the octopaminergic neurons ^[34]^.

In fruit flies, at least two neuropeptides, tachykinin and the short neuropeptide F, are expressed within the CX; their suppression leads, respectively, to increased bout frequency and augmented speed and distances walked ^[27,35]^.

As in the OKR, where specific DNs vehicle the information down to the motor neurons regulating head movements (DN9), the modulation on dopaminergic DNs – with possible circuits to the ventral nerve cord for both the CX and MB ^[36,37]^: *CX → lateral accessory lobes (LAL) → posterior slope (PS); MB → superior medial protocerebrum (SMP) → LAL → PS* – reaching the thorax, could be a good candidate. However, when Tschida and Bhandawat (2015) ^[38]^ studied a couple of such DNs, they found evidence of the correlation between the activity of those DNs and specific patterns of leg movement and walking speed, but they did not notice any difference when stimulating flies with attractive odours eliciting strong responses in terms of leg movements. This observation showed that, at least in these neurons, the major modulation happens during leg movement and may not be influenced by the nature of the stimulus provoking them. Interestingly, recent studies (Feng *et al*., 2024; Rayshubskiy *et al*., 2025) ^[39,40]^ characterized several DNs types (DNa01, DNa02, DNa03) involved specifically in turning control during walking, which nonetheless do have indirect connections with wings’ motor neurons and are engaged in stimulus-directed steering in walking (DNa2) ^[40]^, or have been shown to also promote steering during flight (DNa3) ^[39]^.

The quantitative changes we illustrated here are indeed partly described by an increased average velocity and a reduced interval between bouts (“stop” length), compatible with reduced inhibitory regulation from the MB-CX, which in turn would promote an increased motor activity in presence of the aversive odour. In just one specific case, **E1**, a minor increase in velocity was also associated with a trajectory of decisively opposite direction with respect to the control group. It is possible to assume a more complicated scenario than during a rigidly tethered flight OKR paradigm, which may involve more complex responses to arousal and novelty (see for example **E05** which trajectories are akin to controls while presenting an enhanced motor activity despite experiencing a known, to *D*. *melanogaster*, repellent stimulus) as well as decision-making processes gated through the CX-MB.

One last point worth mentioning is that we did not observe, apparently, any evidence for possible memorization and learning processes in the terms of an enhancement of the response as a function of time, as no major differences in the variables we measured did change over the trials. On the other hand, neither a “decline” in performance was observed. The only exception to this trend was found in the average velocities of **C** and **I05**, which showed a decline over time. Possible explanations may be fatigue or habituation. However, we consider the case of fatigue less likely, as we would have expected still a more widespread decrease in the average speed even in presence of stronger aversive odours, which provoked a change in the direction of movement. It seems therefore likely that perceived aversive odours could have helped keeping the flies engaged during the task, as they actively tried to move away, but did not trigger a – maybe unnecessary – learning process for achieving a “better” escape behaviour as the aversive stimulation kept being around. However, resolving these last speculations will require ad hoc experimental paradigms.

In conclusion, the results we obtained show that the substances commonly used in many publicly available products effectively affect fundamental mechanisms, like the OKR, or intervene in the modulation of more complex processes, as in the OMR, which constitute the basis of a complex repertoire of behaviours concurring to the navigational abilities of insects. As we still have limited knowledge on how and why the final output (behaviour) we observe from this multimodal sensory integration is achieved and selected among other possibilities, the fact that some compounds could act on unpredicted components of behaviours in a plethora of species sharing the same neural architecture and organisation should be worrisome: more study is necessary to explore and better characterize the deeper effects of repellent substance in order to fine tune their application while avoiding any abuse which could become harmful against unwanted or protected targets, such as pollinator insects.

## Materials and methods

### Fly strains

The Berlin-K strain was kindly provided by Prof. Christian Wegener (University of Würzburg).

Flies were reared in vials containing 10 ml of standard cornmeal medium, with a 12h:12h dark-light cycle with controlled temperature (26 °C, at 45 ± 10% humidity). Female flies were separated from males and collected under CO_2_ anaesthesia within 24 h from pupal eclosion, then given at least 24 hours to recover. Adult female flies aged between 3 and 6 days were tested within 6 hours from the light onset.

Every group of flies was evaluated for only one (1) repellent and concentration, meaning that one group experienced, e.g., eugenol, concentrated at either 0.5% or 1%.

Each recording was manually checked before the tracking, and only the flies which walked consistently throughout the duration of the experiments were tracked. Likewise, bad tracking results, which could be caused by illumination issues and/or excessive self-grooming by the flies, were discarded altogether and are not part of the analysis shown in this paper.

### Fly preparation

Flies were transferred from the rearing vial to an empty one which was then put on ice. After the cold anesthetization, single flies (one at a time) were transferred to the mounting block, which could be kept cold (down to +4° C) via a Peltier platform laid on a fan heat-sink and carefully placed upright inside the dedicated groove of the mounting support. The Peltier temperature was set to 14° C to minimize the cold experienced by the flies. The tip of a 34-gauge dispensing needle (BSTEAN, Shenzhen Hemasi E-Commerce Co., Ltd., PRC) was dipped in UV hard resin (DecorRom, Shenzhenshi Baishifuyou Trading Co., Ltd., PRC) removing the excess quantity (barely one drop remaining on the tip). The needle was then placed on a support angled at 120° with respect to the fly’s horizontal body axis (considering the frontal portion of the thorax as the 0° angle) and, with the aid of a micromanipulator, lowered onto the fly, touching the centre of the thorax; the resin was then cured for 45 - 60 seconds with a UV torchlight to glue the animal to the pin. The whole tethering procedure took about 2 minutes. Flies were let to recover from the procedure for about 15 minutes while resting on a piece of paper of appropriate dimensions. The flies were then transferred and mounted inside the experimental setup, and we verified that they could easily move while on the Styrofoam sphere. Badly glued or unwilling flies were discarded.

### Experimental apparatus

The experiment was conducted in a home-built dark chamber (components bought from Thorlabs Inc., US), with side access, containing all the hardware.

A support, mounted on a micromanipulator, ends with a syringe attachment where the pin together with the glued fly can be secured, by the side of the monochrome camera (MQ003MG-CM, Ximea GmbH, Germany). The camera is equipped with an infra-red (IR) bandpass filter and two IR (850 nm) LEDs (M850L3, Thorlabs Inc., US) provide the necessary illumination. The camera resolution was VGA 0.3 MP 648 × 488 pixels, with a pixel size of 7.4 μm and a maximum frame rate of 500 frames per second.

The other components of the apparatus include:

- the projector (Lightcrafter 4500, Texas Instruments Inc.), placed in front of the screen, with a refresh rate of 60 Hz, with a maximal output illuminance of 621.5 lux at the centre of the screen and 435 lux at 45° of azimuthal deflection.
- an adjustable, curved, hand-crafted screen (radius = 6.5 cm, height = 13 cm), made of parchment paper, placed in front of the animal (distance of 5 cm), with an illuminance attenuation factor of 10. The actual projected display had an azimuth of ± 90°, an elevation and depression angle of 22.5°, and a resolution of 1280 x 800 pixels.
- the custom walking machinery, composed of the 3D-printed, M-shaped, sphere holder, a Styrofoam ball hand-painted with an irregular black spot pattern, and the air jet pump (ADD HERE) to keep the sphere slightly floating.
- the custom odour-<u>delivery</u> system made up of plastic tubing of various diameters, assorted luers (Ark-Plas Products Inc., US), glass capillaries (GB150F-10, Science Products GmbH, Germany), an Arduino (UNO REV3, Arduino, US) controlled solenoid valve (SIRAI Elettromeccanica S.r.l., Italy), two glass vials containing the solutions, and an air pump (Air Professional 150, PRO.D.AC INTERNATIONAL S.r.l., Italy) for the vaporizing.
- the custom odour-<u>recycling</u> system, consisting of plastic tubing, an externally alimented suction unit (VN-C4 vacuum pump, You Cheng Industrial Co., Ltd., Taiwan), plus the glass flask used to achieve negative pressure and the subsequent suction.

The tubing was placed perpendicularly to the fly, at around 1 cm and 2 cm for the delivery (on the fly’s right) and the recycling (on the fly’s left) system, respectively. Different (clean) tubing, glass capillaries, and vials were used for the different compounds.

Differently from our previous work, we used a dedicated computer (Workstation HP Pro Tower 400 G9 PCI: Win 10; Intel 12th Gen. 12500 6 cores 3 GHz processor; 16 GB RAM) both for delivering the stimuli and the recording, which was achieved directly through the Ximea Software GUI. A U3-LV Labjack (Labjack Corporation, US), in tandem with an Arduino Uno (Arduino^®^, US) system, controlled the synchronization between the stimuli delivery and the recording timings of the camera, which analogically received inputs from the Labjack-Arduino system.

We programmed the Arduino Uno to control the relevant camera triggers synchronized to the Python script controlling the presentation of the visual stimuli (designed through the open-source PsychoPy^©^ toolbox, Open Science Tools Ltd., UK), and the modulation of the solenoid valve. Sampling rate of the camera was locked at 200 frames per second.

### Repellent compounds

The chosen repellent substances (eugenol, lemongrass oil, picaridin, and IR3535^®^ *alias* ‘Nb[n-N-butyl-N-acetyl] aminopropionic acid ethyl ester’) were purchased at the highest purity available (min. 95 %) from Biosynth^®^ Ltd, UK.

### Pilot study experimental paradigm

The paradigm is structured similarly as described in the further section (seen Suppl. Fig. 1 for a graphical representation). This paradigm derives from a former version we utilized in a previous study where the two MOP (Mineral oil only Phase) and OP (Odorant Phase) phases run consecutively (30 sec. each) with no pause in between.

### Experimental paradigm

The paradigm was structured in 2 (two) quick “training” phases to get flies acquainted to their tethered condition, followed by 4 (four) repetitions of the proper trial, which was subdivided in 2 portions (Fig. 1a). Within the training, as well as during the first part of the trial, flies faced visual stimulation (moving in the same direction throughout all the experiment; same as in the second part) without experiencing any odour, as they were instead delivered a plume coming from a plain mineral oil solution (Mineral Oil Phase, MOP). Within the second part of the same trial, the solenoid valve switched at the set time, causing the air plume to come instead from the vial containing the repellent-enriched mineral oil solution (Odorant Phase, OP). The training and trial were structured as follows:

#### Training

a. **SF**: one 12° vertical black bar on a white background, presented in the middle of the screen (duration: 5 seconds).
b. **MOP**: a mask of gratings made up of 12° alternate black and white vertical bars (spatial wavelength of 24°) moving clockwise at a fixed speed of 15 deg·s^−1^ for a duration of 60 seconds.
c. Same as (*a.*).
d. Same as (*b*).
e. **Pause** in darkness (duration: 20 seconds).

#### Trial

a. **SF**: one 12° vertical black bar on a white background, presented in the middle of the screen (duration: 10 seconds).
b. **MOP + OP**: a mask of gratings made up of 12° alternate black and white vertical bars (spatial wavelength of 24°) moving clockwise at a fixed speed of 15 deg·s^−1^ for a duration of 60 seconds.
c. **Pause** in darkness (duration: 20 seconds).

Stimuli were drawn at the maximum contrast available, resulting in a Michelson contrast of 0.61.

For further clarification, the air plume was continuous, from the start of the training to the end of the trials and was coming always from the same direction, from the right to the left of the animal, which was the direction opposite to the optokinetic stimulation. The Control group never experienced the OP, but two consecutive MOPs instead.

### Data extraction

Recordings were manually checked and only flies which kept an active walking behaviour (i.e., moving > 90% of the time during the optomotor task) were kept and analysed.

The tracking of the animal was conducted offline through the self-contained FicTrac^[41]^ program: we manually set from the GUI, for each recording, the sphere’s ROI and the masks for identifying the fly and the pins (Fig. 1b); further adjustments were made to the settings file for the binarization and the thresholding of the input video, as suggested by the FicTrac’s author. The recording was checked live to assess the correctness of the automated tracking procedure through the debugging FicTrac feature.

The FicTrac output data (X, Y, Z coordinates, and the timestamps) were imported into RStudio for the subsequent analysis (Fig. 1c).

X, Y, and Z data were smoothed applying a simple moving average (SMA) with 10 *sample length* before further calculations of the related per-fly weighted means. Mean velocities were computed as the total path length (sum of the position’s differentials absolute values in degrees) walked over 60 seconds (trial duration). Instant velocities and accelerations were not directly imported from the FicTrac output, but rather manually computed after smoothing the position data with a Savitzky–Golay low-pass filter with a cut-off at 10 Hz to remove high frequency noise.

Regarding the analysis on the “*stop*” and “*go*” windows, we defined as “*stops*” the time windows where 𝛼_ins_ < 10% of the time-averaged 𝛼_ins_. Equally, “*go*” were defined by values higher than the set 10% threshold, plus we only considered periods as proper “*go*” phases those lasting at least 100 milliseconds. Then, we tagged the average *α*_ins_ values as coherent (+) or non-coherent (-) over the minimum span of *bona fide* “go” periods we set (100 ms), when the ratio of the negative *α*_ins_ versus positive *α*_ins_ showed to be unbalanced towards one of the two (***-***/***+***) *α*_ins_ with a minimum threshold of 60%.

### Statistical analysis

The whole analysis was conducted in R-Studio (v4.4.1.) with a significance level *α* = 0.05. Data were checked for normality (Shapiro-Wilkins test) and heteroscedasticity (Bartlett’s test when the distributions were not normal) across groups.

In the pilot experiment, statistical significance regarding velocity and rotation amount were assessed through ANOVA + Tukey HSD. The differences between trajectories were analysed by comparing the slopes of the regression lines, obtained from the ‘*lm*’ function, with the ‘*emtrends*’ and ‘*pairs*’ functions (EMMs, “emmeans” R package).

In regard to the main experiment, the statistical significance for trajectories and quantity of movement (not all groups had normal distributions, and the variance was no homogeneous) was assessed through Kruskal-Wallis plus post-hoc Dunnet test and Pairwise Test for Equality of Proportions respectively, both adjusted with Benjamin-Hochberg correction for multiple comparisons – which we preferred to the Bonferroni correction (used later) due to the multiple levels in the test. We also run Wilcoxon signed rank exact test comparison within each group to compare the result with the multifactorial Kruskal Wallis + Dunnet test, but we did not find significant differences between the “MOP/OP” phases.

In regarding to the analysis of the mean velocities, to reduce the effect of outliers, we compared the results from the Kruskal-Wallis and post-hoc Dunnet test with robust ANOVA for trimmed means followed by the related post-hoc mcppb20 test with Bonferroni adjustment (“WSR2” R package), which we refer to in the results as it proved more conservative.

Not all groups had their instantaneous velocities values normally distributed; therefore, for a multi-factorial analysis of velocities and “Stop” and “Go” periods we opted for a non-parametric ANOVA after aligned rank transformation test (“ARTool” R package) for assessing the relevant effects and interactions to then build a linear model for subsequent Estimated Marginal Means (EMMs, “emmeans” R package) computation and pairwise comparisons with Bonferroni adjustment.

## Abbreviations

C: control group
CX: central complex
E05: group exposed to eugenol 0.5% solution
E1: group exposed to eugenol 1% solution
HOKN: head optokinetic nystagmus
I05: group exposed to IR3535^®^ 0.5% solution
I1: group exposed to IR3535^®^ 1% solution
L05: group exposed to lemongrass 0.5% solution
L1: group exposed to lemongrass 1% solution
IR: infra-red
MB: mushroom bodies
MOP: mineral oil phase
OF: optic flow
OKR: optokinetic response
OMR: optomotor response
OP: odorant phase
P05: group exposed to picaridin 0.5% solution
P1: group exposed to picaridin 1% solution
SF: stripe fixation

## Data Availability

An R-markdown file retracing the content of the paper is available in the “OMR-Repellents” repository ([Github](https://github.com/Aramicus/OMR-Repellents.git)), where a google-drive link is also provided to access the original generated Dataframes. Additionally, the pre-processed data, along with a video sample are hosted in the “*Repellent olfactory cues influence the optomotor response modulation in Drosophila melanogaster - Data*” repository at Zenodo.com (DOI: https://doi.org/10.5281/zenodo.18416157).

## Author contributions

GMM, PV, AD, and AM designed the experiment. GMM, MB, and AM set up the experimental apparatus and wrote the related scripts. GMM designed both the experimental paradigms. GMM and SZ acquired the data. GMM carried out the pre-processing (together with SZ) and the data analysis. GMM drafted the manuscript, with contribution from MB, AM, and MDM, and prepared the figures and tables. All authors contributed to the interpretation of data and the revision of the manuscript.

## Acknowledgements

Authors thank Prof. Christian Wegener, from Biocenter - University of Würzburg (DE), who kindly provided the Berlin-K *Drosophila* strain, and Dr. Paola Cisotto, from the Biology Department - University of Padova (IT), who supported us with the caring and nurturing of the flies’ strains.

